# Concentration-dependent reduction of planktonic- and biofilm-state *Vibrio alginolyticus* by the bacteriophage pVa-21

**DOI:** 10.1101/322933

**Authors:** Sang Guen Kim, Sib Sankar Giri, Jin Woo Jun, Saekil Yun, Hyoun Joong Kim, Sang Wha Kim, Jeong Woo Kang, Se Jin Han, Dalsang Jeong, Se Chang Park

## Abstract

There is an increasing emergence of antibiotic-resistant *Vibrio alginolyticus*, a zoonotic pathogen that causes mass mortality in aquatic animals as well as human infection; therefore, there is a demand for alternatives to antibiotics for treatment and prevention of infections caused by this pathogen. One possibility is through the exploitation of bacteriophages. In the present study, the bacteriophage pVa-21 belonging to *Myoviridae*, was isolated and characterized as a candidate biocontrol agent against *V. alginolyticus*. Its morphology, host range and infectivity, growth characteristics, planktonic or biofilm lytic property, stability under various conditions, and genome were investigated. Its latent period and burst size were estimated to be approximately 70 min and 58 plaque-forming units/cell, respectively. In addition, phage pVa-21 could inhibit bacterial growth both in the planktonic and biofilm state. Furthermore, phylogenetic and genome analyses revealed that the phage is closely related to ‘phiKZ-like phages’ and can be classified as a new member of the phiKZ-like phages that infect bacteria belonging to the family Vibrionaceae.

## Introduction

*Vibrio alginolyticus* is an opportunistic pathogen frequently found in marine environments. Having a wide host range, it can cause mass mortality in aquatic animals as well as human infection [1–4]. It is well known that the majority of bacterial infections are caused by bacteria in biofilms [5]; therefore, biofilm formation is an important feature of pathogenic microorganisms [6]. Furthermore the threat of biofilms is apparent from the fact that bacterial cells in biofilms are highly tolerant to antibiotics compared with those in the planktonic state [7]. Therefore, antibiotics cannot effectively inhibit bacterial proliferation within biofilms resulting in regrown bacteria following failure of antibiotic treatment, which could cause serious problems [8]. Therefore, there is a need for effective alternatives to antibiotic treatment for managing biofilm-related bacterial outbreaks.

As viruses of bacteria, bacteriophages (phages) specifically infect and lyse the targeted bacteria. Recently, lytic bacteriophages have been demonstrated as potential alternatives to antibiotics in various studies targeting the prophylaxis or treatment of bacterial diseases [9–11]. Especially in case of biofilm-related outbreaks, phage-infected bacteria existing at the outermost region of the matrix can play a pivotal role in further spreading the phages into a biofilm complex. The amplification of bacteriophages within the biofilm and the enzymes they produce have been studied in various bacterial biofilms [12, 13].

In theory, bacteria, whether in the planktonic or biofilm state, can be degraded by a single phage particle as its progeny infect and disrupt the adjacent cells. Previous studies have suggested that phages can control biofilms effectively [14]. However, low phage concentrations allow bacterial growth. Therefore, for practical applications, it is necessary to investigate and determine the minimum inhibitory concentration for phages as in the case of antibiotics. The present study was designed to establish the importance of the minimum bactericidal concentration (MBC) of phages against bacterial cells both in the planktonic and biofilm states.

In the present study, with the goal of improving the treatment and prevention of *V. alginolyticus* infection, we isolated and characterized a new lytic phage (designated as pVa-21) infecting *Vibrio* strains, which is part of the harveyi clade and a major aquatic animal pathogen. The biological characteristics of the *Myoviridae* phage pVa-21 were evaluated and its complete genome was sequenced and analyzed. Characterizations were focused on both the anti-planktonic and anti-biofilm state cell activity of the phage.

## Materials and Methods

### Bacterial strains and growth conditions

All the bacterial strains used in this study are listed in S1 Table. The bacterial strains were cultured in tryptic soy broth (TSB; BD, Sparks, MD, USA) supplemented with 1.5% (w/v) of sodium chloride (Daejung Chemicals, Korea) or sub-cultured on tryptic soy agar (BD).

### Phage isolation, purification, and propagation

To isolate phages infecting *V. alginolyticus*, water samples from the West Sea of Korea were collected. Water was filtered through 0.2-μm membrane filters (Merck Millipore, Billerica, MA, USA). Strain rm-8402, which was previously reported as a fish pathogen [2], was used as an indicator host strain. To isolate the phage, strain rm-8402 was co-cultured with the collected water samples for 24 h at 27°C. After enrichment, the culture was centrifuged at 10,000 × *g* for 20 min and the resulting supernatant was filtered through a 0.2-μm membrane filter. To verify the presence of the lytic phage in the filtrate, the double-layer agar method was performed using the filtrate [15]. After overnight incubation at 27°C, the plaque was purified five times through single-plaque isolation with a sterile straw to ensure that the isolated phage represented a descendant from a single virion.

### Electron microscopy

The obtained phages were concentrated using polyethylene glycol 8000-NaCl precipitation in SM buffer [100 mM NaCl, 50 mM Tris (pH 7.5), and 10 mM MgSO_4_], and 10 μ1 of the suspension was spotted on a copper grid. After 2 min, the suspension was removed by absorption onto filter paper and the phages were negatively stained with 2% uranyl acetate for 1 min, followed by three successive washes with water. The grid was air-dried for 10 min and observed with a JEM-1010 (JEOL, Tokyo, Japan) transmission electron microscope (TEM) operated at 80 kV. The dimensions of phages were calculated by measuring five independent phages.

### Host range analysis

The host range of the obtained phage (designated pVa-21) was determined using a spot assay. Ten microliters of the phage lysate [>10^7^ plaque-forming units (PFU)/ml] was dropped onto the overlaid top agar and mixed with each bacterial strain. The plates were then incubated overnight at 27°C and checked for the presence of plaques. An efficiency of plating (EOP) assay was conducted to quantify the lytic activity of phage pVa-21. The phage suspension (10^3^ PFU/ml) was assayed by the double layer agar method. The total number of plaques was determined after 24 h incubation and EOP values were calculated by comparing the ratios of PFU of a susceptible strain to the indicator strain rm-8402 in triplicate.

### Adsorption assay and one-step growth curve

The adsorption assay was carried out as described by Lu et al. (2003). The exponentially growing host strain [1.5 × 10^8^ colony-forming units (CFU)/ml] was infected with a phage suspension at a multiplicity of infection (MOI; the ratio of virus to bacterial cells) of 0.001 and incubated at 27°C. Aliquots of 100-μ1 were taken at 0, 5, 10, 15, 20, 25, and 30 min after infection and immediately diluted in 900-μ1 of PBS and centrifuged at 12,000 × *g* for 5 min. The supernatants were titrated for un-adsorbed free phages using the double-layer agar method. To construct the growth curve, the phage lysate was inoculated in 10 ml of exponentially growing host strain culture (1.5 ×10^8^ CFU/ml) at an MOI of 0.001. The phage was absorbed for 15 min and then centrifuged at 12,000 × *g* for 5 min. After the supernatant was discarded, the phage-infected bacterial pellet was re-suspended in 10 ml of preheated TSB and incubated at 27°C with shaking at 250 rpm. At 10-min intervals, 100-μ1 aliquots were taken until 140 min, and the titer was immediately determined by the double-layer agar method. The titer measurements were carried out in triplicate.

### pH and thermal stability assessment

For pH stability tests, the phage suspension was inoculated in 1 ml of phosphate-buffered saline (PBS)-containing tubes adjusted to pH 3.0, 5.0, 7.0, 9.0, and 11.0 with 1 M NaOH or 1 M HCl. The tubes were then incubated at 27°C and aliquots were taken at 60 min. For thermal stability tests, the phage suspension was incubated at 4°C, 20°C, 25°C, 30°C, 35°C, 40°C, and 50°C, and aliquots were collected at 60 min. All tests were performed in triplicate.

### Planktonic cell treatment with pVa-21

To evaluate the bacteriolytic efficacy of pVa-21, strains showing a turbid or clear lysis pattern in the spot assay were selected. One percent of overnight culture was inoculated into 10 ml of fresh broth to obtain 10^8^ CFU/ml, and the phage was inoculated at a MOI of 0, 0.1, 1, and 10. Planktonic cells were cultured with vigorous shaking and optical density at 600 nm (OD_600_) was measured at 0, 1, 3, 5, 7, 9, 12, and 24 h. All tests were performed in triplicate.

### Biofilm treatment with pVa-21

To verify the anti-biofilm efficacy of pVa-21, a biofilm was formed according to a previously described method with minor modifications [17]. In brief, the biofilm assay was performed in 96-well polystyrene tissue culture microplates (Nunc, Roskilde, Denmark). One percent of overnight culture was inoculated into fresh TSB supplemented with 1% D-glucose (Sigma Aldrich, St. Louis, MO, USA) and aliquots of 200 μ1 were distributed in each microplate well. The microplate was then incubated at 27°C for 24 h with no shaking. At 24 h, the supernatant of each well was removed and washed twice with PBS to remove all planktonic cells, and then treated with 200 μl of phage suspension at a series of concentrations, low (10^5^ PFU/ml), middle (10^7^ PFU/ml), and high (10^9^ PFU/ml), for 24 h. The microplate was washed twice with PBS and dried sufficiently. The formed biofilm was stained with 1% crystal violet to quantify the total biomass and its optical density at 595 nm was measured.

### Sequencing and genome analysis of the phage

The genomic DNA of the phage was extracted as described previously [18]. The DNA of pVa-21 was then digested with 10 U of DNase I, and RNase A according to the manufacturer’s protocol. The purified genomic DNA of the phage was sequenced using an Illumina Hiseq 2500 platform at Genotech (Daejeon, Korea). Reads were trimmed and assembled using the CLC Genomic Workbench v6.5.1. Putative open reading frames (ORFs) were predicted and annotated using Glimmer v3.02 [19], Prodigal v1.20 [20], and protein BLAST, respectively. The Rapid Annotation using Subsystem Technology (RAST) server was used for confirmation [21]. Detection of tRNAs was carried out using tRNAscan-SE v2.0 [22], and nucleotide homology was determined using EMBOSS Stretcher [23]. The genome map of pVa-21 was drawn using DNA plotter [24]. The web tool RESFINDER v2.1 was used to search for known antimicrobial resistance coding genes [25]. The amino acid sequence of the major capsid protein and terminase large subunit was obtained from Genbank database and was aligned using Clustal W [26]. A phylogenetic tree was constructed using the maximum likelihood method implemented in MEGA version 7.0 [27] with 1000 bootstrap replications.

### Nucleotide sequence accession numbers

The genome sequence of the isolated phage pVa-21 was deposited in GenBank under the accession number KY499642.

### Data analysis

All analyses were performed using SigmaPlot 12.0 software (Systat Software, Inc. Chicago, USA).

Comparisons between different concentration points and positive controls were conducted by performing Student’s t-test. A value of P < 0.05 was considered statistically significant.

## Results

### Isolation and biological properties of phage pVa-21

One *V. alginolyticus* phage was isolated from sea water samples after enrichment. The purified phage was examined by TEM and classified based on the criteria proposed by Ackermann (28). As shown in Fig 1, pVa-21 was assigned to the family *Myoviridae* with an icosahedral head of 87 ± 3 nm in diameter (n = 5) and a contractile tail of 240 ± 9 nm in length (n = 5). The host range test was determined against bacteria of the Harveyi clade, which include major pathogens of aquatic organisms, including *V. alginolyticus* (n = 5), *V. harveyi* (n = 5), *V. parahaemolyticus* (n = 1), *V. anguillarum* (n=1), *V. campbellii* (n = 1), and *V. vulnificus* (n=1). Phage pVa-21 was able to infect *V. alginolyticus* (n = 3) as well as *V. harveyi* (n = 1) resulting in a turbid or clear plaque pattern (S1 Table). However, the phage did not show infectivity to the nine other bacterial strains tested. Adsorption rate was assessed as described above, and the percentages of adsorbed phages in the *V. alginolyticus* rm-8402 after 15 min was 95% (Fig 2). To identify the growth pattern and burst size of pVa-21, a one-step growth curve was generated. The latent period was found to be approximately 70 min and the burst size, i.e., the number of progeny released after the lysis of a single bacterial cell, was approximately 58 virions per cell (Fig 3). The stability of pVa-21 was tested at various pH and temperature conditions and was estimated by determining the changes in growth (based on the number of PFU). Phage pVa-21 showed relatively good stability at pH 5.0, 7.0, and 9.0 after 1 h of incubation, but significant decreases in PFU counts were detected at low and high pH. At pH 3.0 and 11.0, pVa-21 was extremely unstable with a nearly 100% decrease in the PFU observed (S1 Fig A). The thermal stability test showed that pVa-21 was quite stable (> 90%) at 4°C, 20°C, 25°C, 30°C, and 35°C for 1 h, but its numbers sharply decreased at temperatures above 40°C (S1 Fig B).

**Fig 1.**
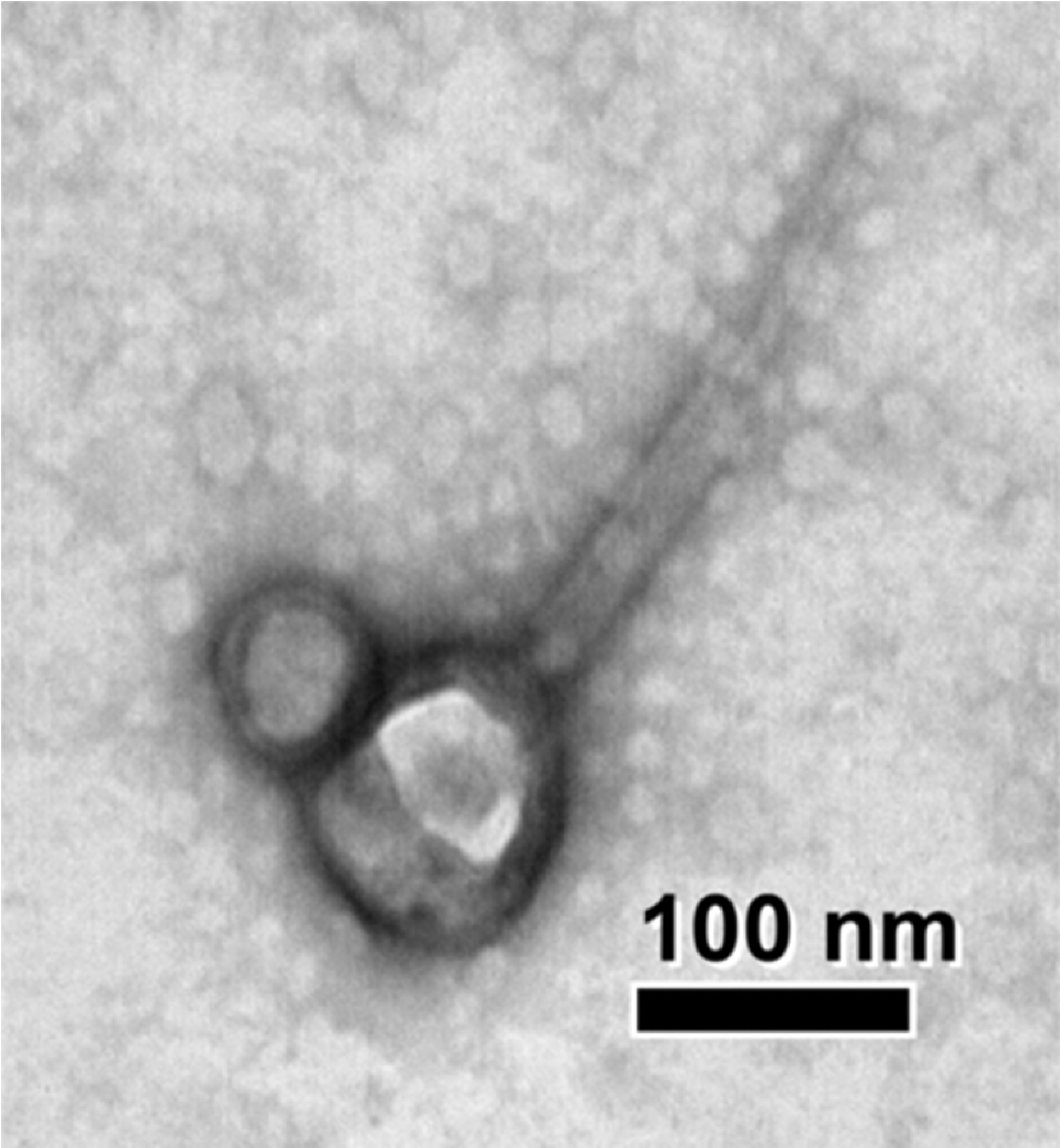
Transmission electron micrograph of pVa-21. The virion was negatively stained (bar = 100nm).

**Fig 2.**
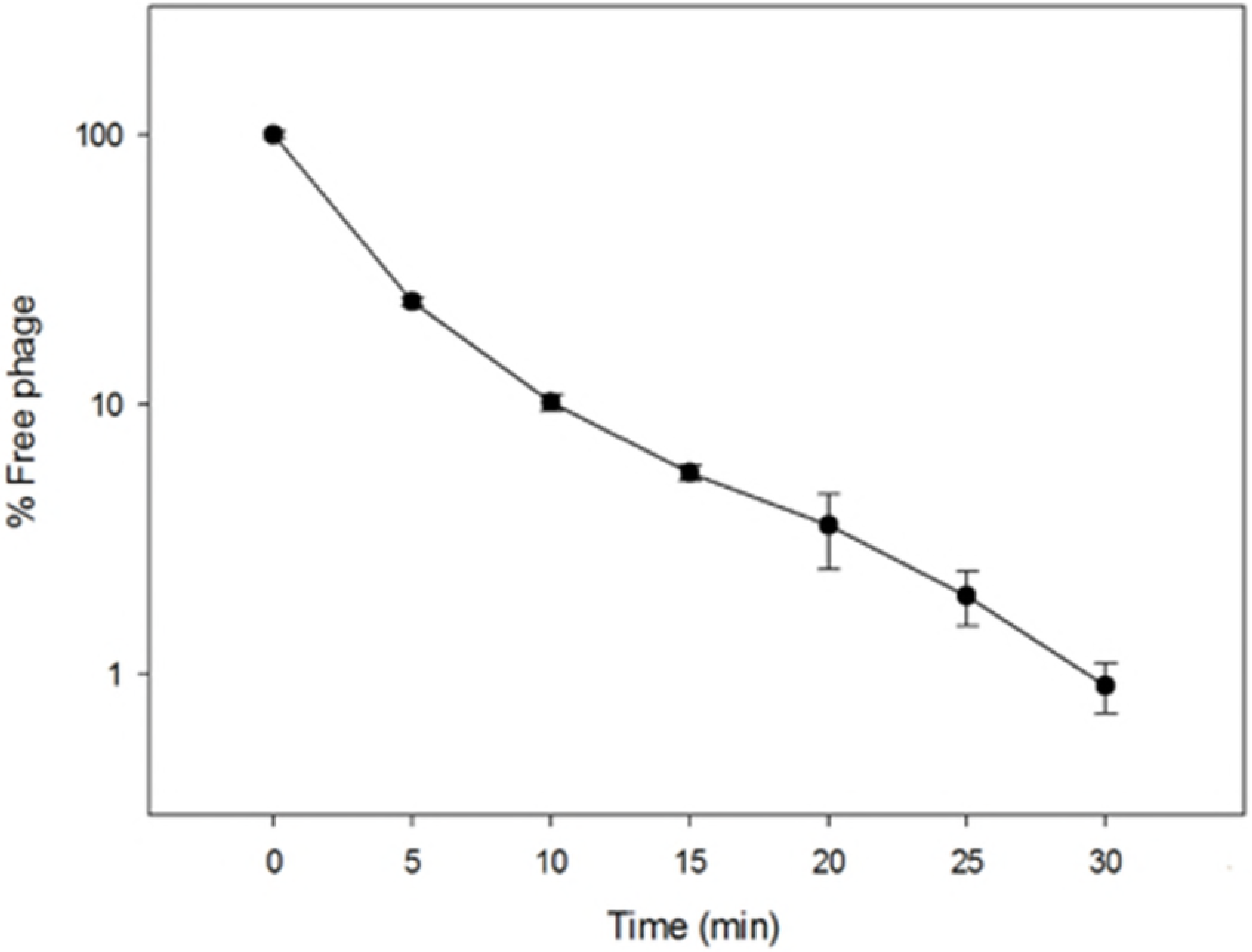
Adsorption of pVa-21. Phage pVa-21 was added to exponentially growing *V. alginolyticus* rm-8402 strains. The results are shown as mean ± standard deviations from triplicate experiments.

**Fig 3.**
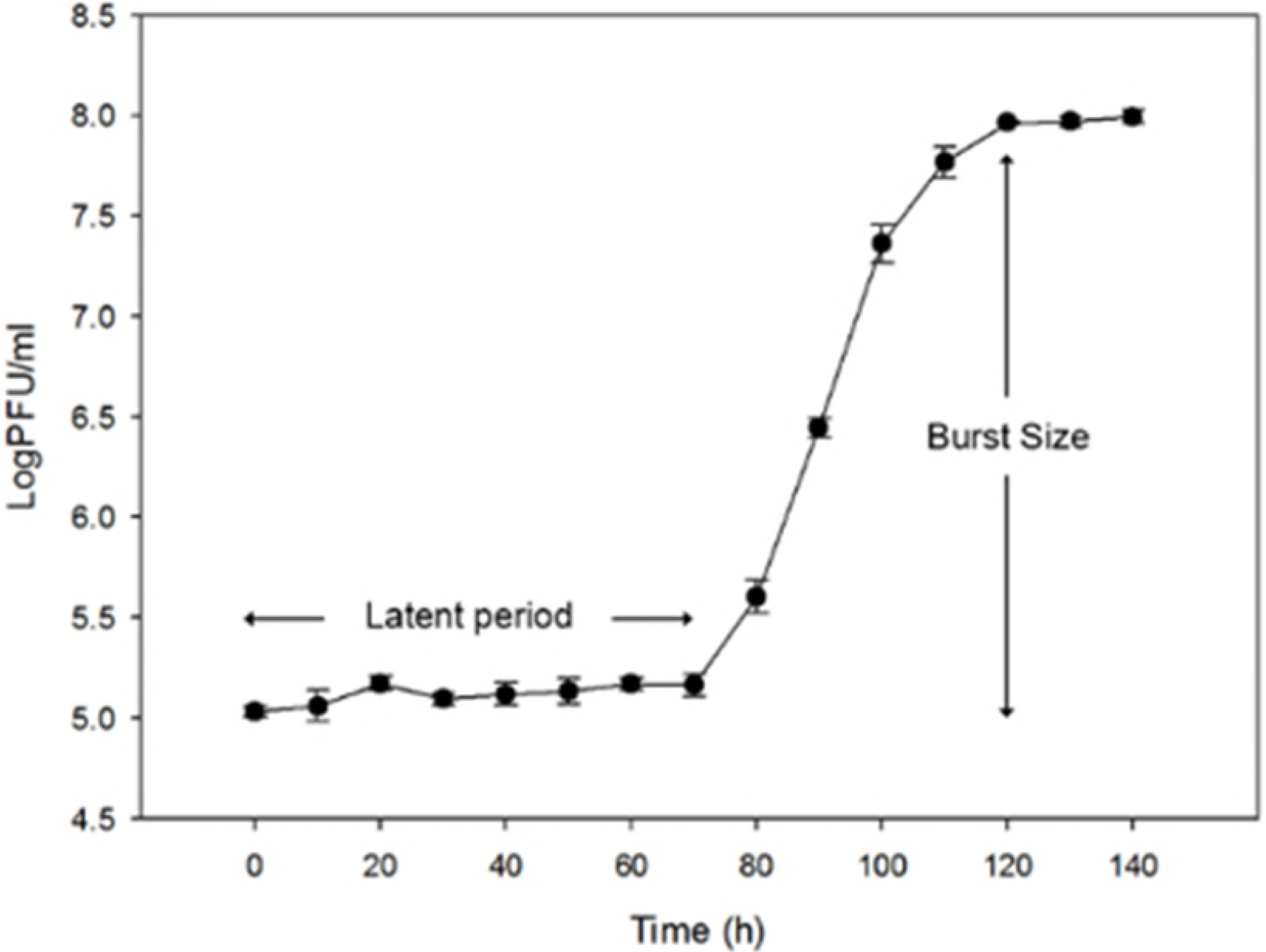
One-step growth curve of phage pVa-21 in *V. alginolyticus* rm-8402 strain. The results show mean ± standard deviations. Latent time and burst size were inferred from triplicate experiments.

### Planktonic and biofilm cell treatment with phage pVa-21

Among the 18 *Vibrio* strains used in present study, the 10 strains showing a turbid or clear lysis pattern were used for the planktonic cell treatment test. Strains that were not inoculated with phage pVa-21 (MOI: 0) showed a continuous increase in the OD600 value for 24 h of incubation. In contrast, growth of the reference strain rm-8402 increased until 3 h and was then completely inhibited by the phage at an MOI of 10 and 1 (Fig 4A). Strains V447 and SFC-BS showed a turbid lysis pattern in the spot assay, and were inhibited at an MOI of 10, reaching an OD_600_ value of almost 0 after 7 h of incubation. Although the growth of these two strains was also affected by the phage at an MOI of 1, the populations were sustained to some extent and did not reach a value of 0 (Fig 4B and D). Similarly, the growth of strain am-10 was affected by pVa-21 in proportion to its inoculated concentration (Fig 4C). The remaining strains did not appear to be affected by phage pVa-21 showed no differences in growth compared to the respective un-inoculated (MOI 0) group. When the lowest concentration of phage, MOI 0.1, was treated, all strains showed no growth inhibition by the phage except strain rm-8402. The three strains that reached an OD600 value of 0 in the planktonic cell treatment assay were further subjected to the biofilm treatment assay. The 24-h-old biofilm formed by each of the three strains was challenged with low (10^5^ PFU/ml), middle (10^7^ PFU/ml), and high (10^9^ PFU/ml) concentrations of the phage for 24 h, except for strain SFC-BS as it could not form a biofilm over 24 h or until 48 h (S2 Fig). The biofilm of strain rm-8402 was significantly disrupted (P < 0.05) when treated with the middle or high phage concentration, but showed no significant decrease upon treatment with the low concentration of pVa-21 (Fig 5). The phage more effectively disrupted the biofilm of strain V447, resulting in an OD value less than 2 (P < 0.05) at the low concentration, with further disruption to an OD value of ≤ 1 (P < 0.05) at the middle or high concentration (Fig 5).

**Fig 4.**
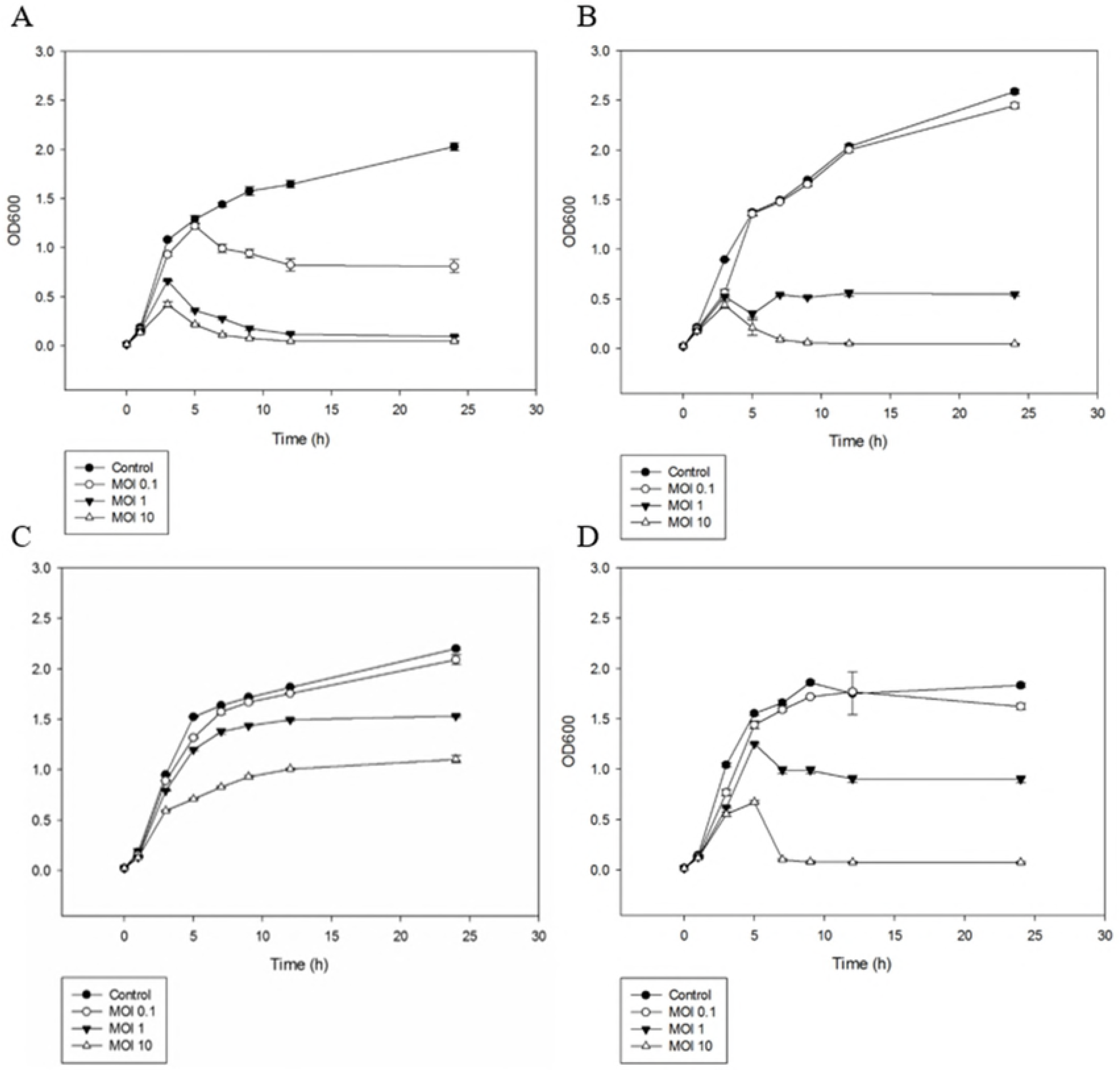
The planktonic cell lytic activity of pVa-21 against 4 strains of *Vibrio* species. Overnight cultures of *V. alginolyticus* rm-8402 (A), *V. alginolyticus* V447 (B), *V. alginolyticus* am-10 (C), and *V. harveyi* SFC-BS (D) were co-cultured with pVa-21 at MOI of 0, 0.1, 1, and 10. The results are shown as mean ± standard deviations from triplicate experiments.

**Fig 5.**
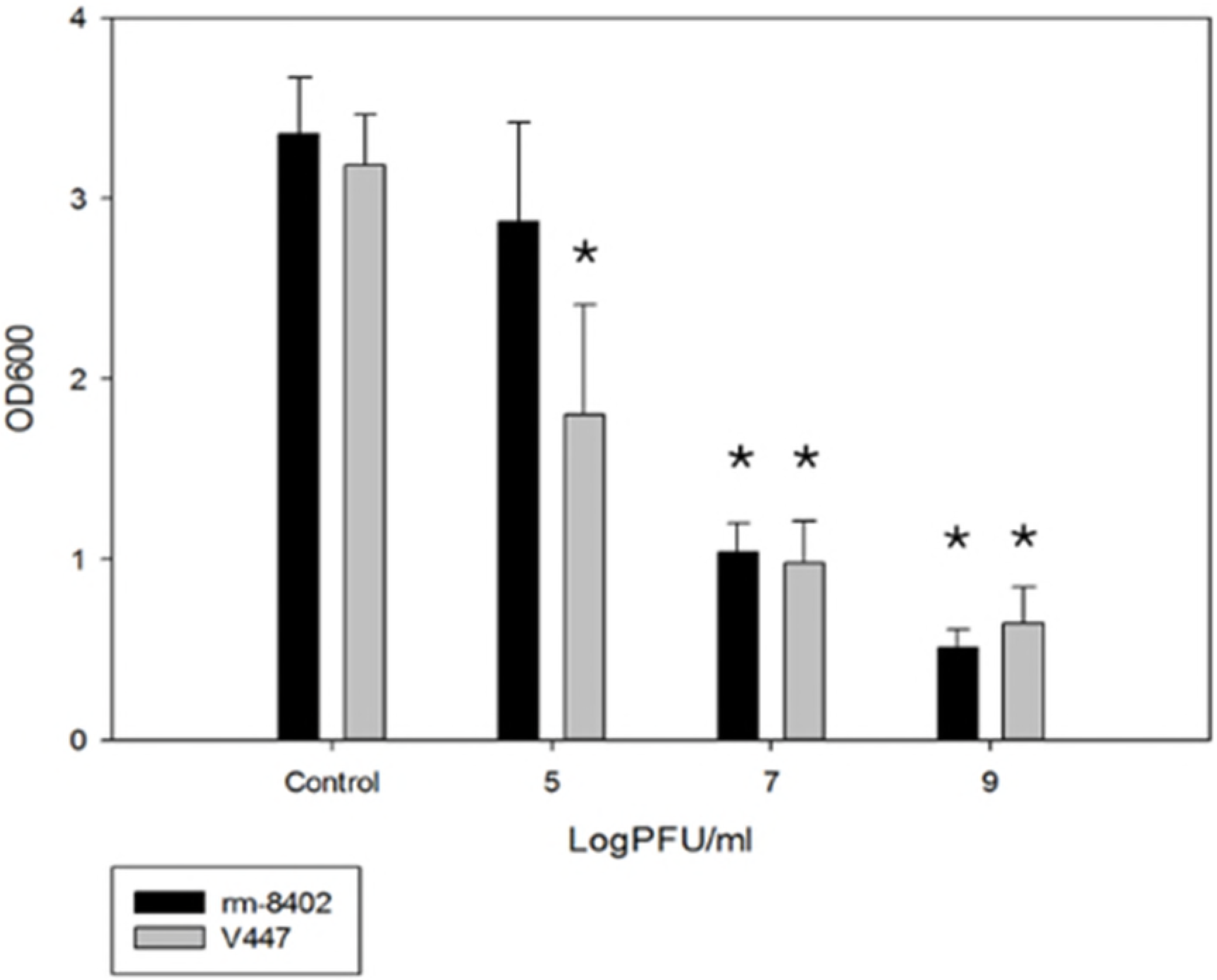
The biofilm lytic activity of pVa-21 against 2 *Vibrio alginolyticus* strains. Biofilms (24 h old) of *V. alginolyticus* rm-8402 and *V. alginolyticus* V447 were treated with pVa-21 at concentrations of 10^5^, 10^7^, and 10^9^ PFU/ml. The results are shown as mean ± standard deviations from triplicate experiments.

### Genomic characterization of phage pVa-21

The whole genome of phage pVa-21 was sequenced and analyzed. In general, the genome of phages belonging to the family *Myoviridae* is known to consist of double-stranded DNA (dsDNA). In line with this common observation, the genomic DNA of pVa-21 was digested by DNase I and not by RNase A suggesting that its genome comprised DNA. The complete genome sequence of the linear phage pVa-21 consists of 231,998 bp with a GC content of 44.58%, encoding 241 putative ORFs and no tRNA genes. As shown in Fig 6, the ORFs of pVa-21 were broadly scattered across the genome and were not clustered according to function, such as those encoding structural, metabolism-related, or lysis proteins. Functional examination of the predicted ORFs showed that they could be classified into three main categories: nucleotide regulation (e.g., helicase, ribonuclease H, DNA-directed RNA polymerase beta subunit), structure and packaging (e.g., major capsid protein, terminase large subunit, tail fiber protein), and lysis (e.g., lytic transglycosylase). The majority of the genes (211, 87.5%) were located in the positive strand with only 30 (13.5%) genes located in the negative strand. Based on its genome size, phage pVa-21 can be classified as a phiKZ-like phage along with the schizo T4-like *Vibrio* phages KVP40, VH7D, and phi-pp2. Moreover, several paralogous genes similar to the beta/beta’ subunit of RNA polymerase were found in the pVa-21 genome, which is a distinctive feature of phiKZ-like phages. The BLASTN search revealed that phage pVa-21 is related to the phiKZ-like phage group (> 65% similarity), including the *Salmonella* phage SPN3US, the *Cronobacter* phage CR5, and the enterobacteria phage SEGD1. As shown in Fig 7, comparison of phage pVa-21 with schizo T4-or phiKZ-like *Vibrio* phages revealed high nucleotide-based homology. The phylogenetic analysis of the major capsid protein and terminase large subunit sequence is shown in Fig 8A and B, respectively. Considering the results of these comparative analyses, phage pVa-21 appears to be more homologous to the phiKZ-like phage group than to the schizo T4-like phage group. Interestingly, three lysis-related proteins were identified in the pVa-21 genome: locus tag pVa21_119 (AQT28060.1), pVa21_134 (AQT28075.1), and pVa21_165 (AQT28106.1). Locus tag pVa21_119, pVa21_134, and pVa21_165 has a lysis catalytic domain similar to the glycosyl hydrolase 108 domain, lytic transglycosylase domain and goose egg white lysozyme domain, respectively. Furthermore, Locus tag pVa21_119 has a C-terminal peptidoglycan-binding domain. S2 Table lists the general features of the putative ORFs identified in pVa-21. The genomic DNA of phage pVa-21 was further examined for the presence of antimicrobial resistance genes, but no such gene was identified.

**Fig 6.**
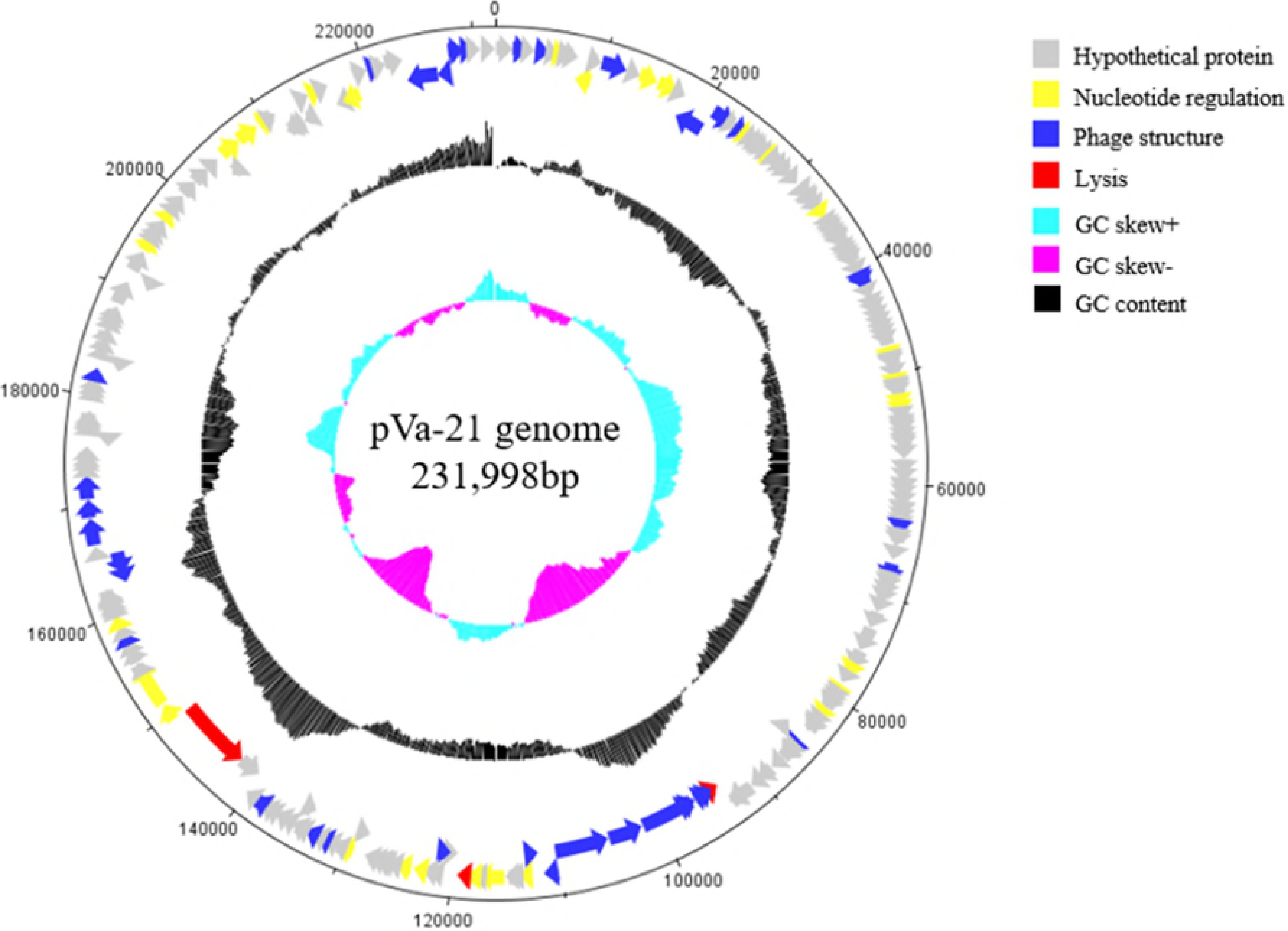
Genome map of phage pVa-21. The innermost circles colored in cyan and purple indicate the GC skew + and GC skew - respectively. Black circle indicates the GC content. The functional categories of ORFs are indicated by specific colors: Grey ORFs represent hypothetical proteins, yellow ORFs represent nucleotide regulation proteins, blue ORFs represent structure and packaging proteins, and red ORFs represent lysis proteins. The scale units are base pairs.

**Fig 7.**
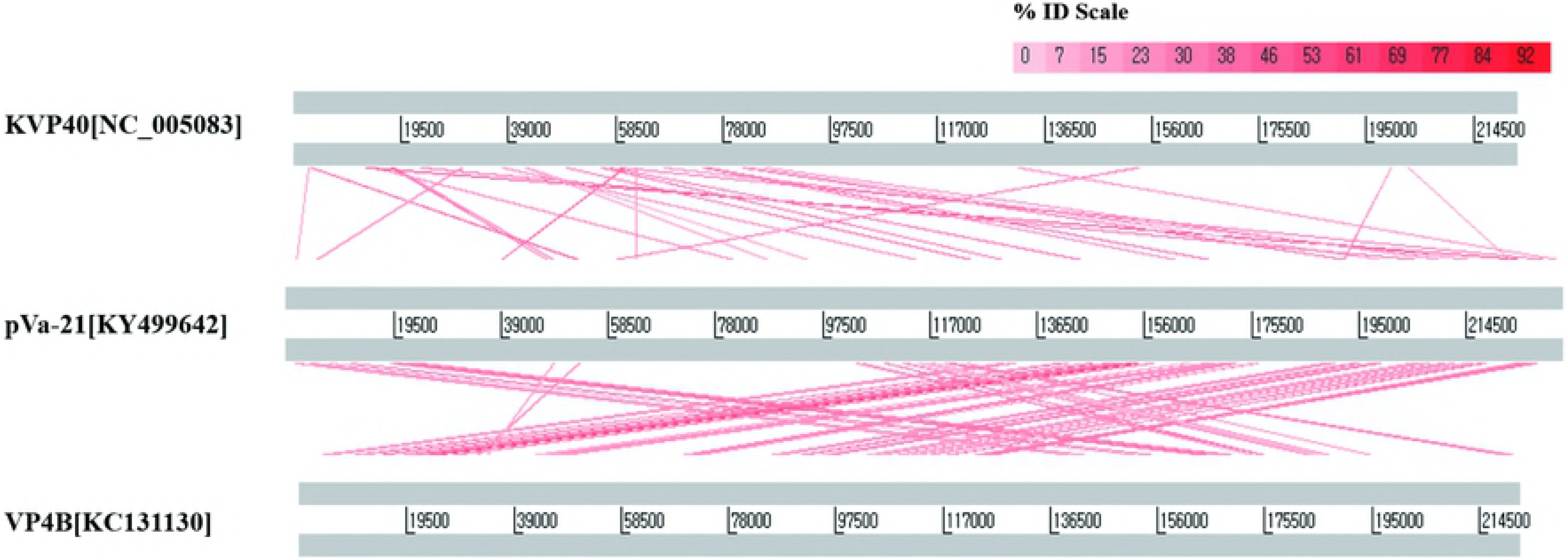
Comparative genome analysis using Artemis comparison tool (ACT). Genomes of KVP40, schizo T4-like phages (top), pVa-21 (middle), and VP4B, phiKZ-like phages (bottom), whose GenBank accession numbers are NC_005083, KY499642, and KC131130, respectively.

**Fig 8.**
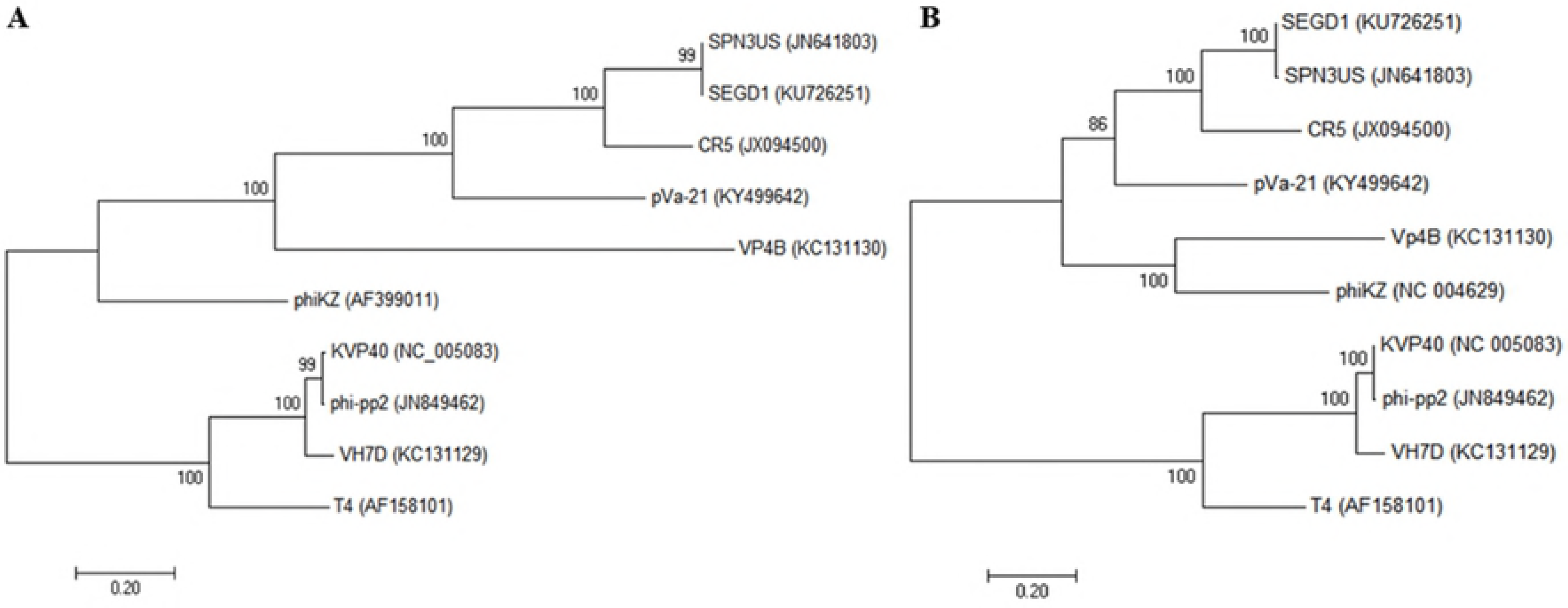
Phylogenetic analysis of various Vibriophages from the T4 or phiKZ phage groups. Major capsid proteins (A), and Terminase large subunits (B) were aligned using ClustalW and a phylogenetic tree was generated in MEGA 7.0 software using the Maximum likelihood method.

## Discussion

It is well known fact that bacteria can grow when the antibiotic concentration is below the minimum concentration. Previously in experiments *in vitro*, lytic phages were also found to be unable to inhibit bacterial growth when their ratio was much lower than bacteria [29], and in phage therapy experiments *in vivo,* the ratio of phages to bacteria, i.e. MOI, was found to be a major discrimination point [30]. Most bacteriophage studies have demonstrated a concentration dependent bacterial cell inhibition effect at planktonic state, but biofilms have received little attention.

In terms of the emergence of resistance bacteria, one of the most important aspects for practical application of phages as antibacterial bio-control agents is to determine the minimum bactericidal concentration to prevent microbes from acquiring resistance to the phage. However, few studies have assessed the difference in the biofilm inhibition effect depending on the initial phage concentration applied [31]. A few studies consider reaction time or the administration number-dependent biofilm eradication effect of phages [32, 33]. With this background, we determined the concentration-dependent bactericidal effect of the isolated phage pVa-21 on bacteria both in the planktonic and biofilm state. In the planktonic state, three strains (rm-8402, V447, and SFC-BS) were clearly lysed by the phage and suffered growth inhibition in a concentration-dependent manner, as shown in other studies [34]. Phage pVa-21 showed a significant anti-biofilm effect by removing the biofilm of each bacteria till the biofilm formed became very thin (OD_595_ ≤ 1). Although bacterial growth was observed in the lower phage concentration group, previous research indicates that regrown bacteria can be controlled by another single inoculation of the phage lysate [32]. Another study suggests that the regrown bacteria recover their susceptibility when subsequently cultured in a phage-free medium and show transient phage tolerance [33].

Several beta/beta’ RNA polymerase subunits are characteristic of phiKZ-like phages [35], and phage pVa-21 had several virion associated beta/beta’ RNA polymerase subunits homologous to those of phiKZ. The phylogenetic analysis shown in Fig. 8 illustrates that phage pVa-21 was clustered more closely to phiKZ-like phages than schizo T4-like phages, and comparative results between pVa-21 and the schizo T4-like *vibrio* bacteriophage KVP40 or the phiKZ-like *vibrio* bacteriophage VP4B support that pVa-21 is more closely related to the phiKZ-like group. Further, morphologically belonging to *Myoviridae,* pVa-21 has a very small plaque size (S3 Fig) which is also one of the characteristics of phiKZ-like phages. No antibiotic resistance or virulence genes were detected in the genome, based on the currently available database, suggesting that the *V. alginolyticus* phage pVa-21 could safely be used as a biocontrol agent against these bacteria. However, the majority of ORFs in pVa-21 did not match with the predicted function in GenBank. Thus, to ensure the safe and reliable application of phages from a therapeutic perspective, further investigation of the phage genome is needed to gain a deeper understanding of the roles of the encoded gene products, as they might produce novel virulence factors or interact undesirably with the host genome [36].

In conclusion, the present study highlights the importance of the initial phage concentration on its antibacterial effects regardless of the growth state of the target bacteria: planktonic or biofilm. The characteristics of the *Vibrio* phage pVa-21 and its genome are expected to broaden the phiKZ-like phage library and help promote the application of bacteriophages in biofilm control to protect aquatic organisms from infection. In addition, as illustrated in a recent study [37], biofilm eradication by phiKZ-like phages in multidrug resistant bacteria suggests that the phiKZ-like phage, pVa-21, could also be used as a bio-control agent. The lytic effect of pVa-21 itself and the three distinct enzymes it possesses are expected to eradicate bacteria. Although the precise biofilm eradication mechanism of pVa-21 remains to be elucidated, the three different lytic enzymes detected are likely involved in the mechanism.

## Acknowledgements

This research was supported by the Basic Science Research Program through the National Research Foundation of Korea (NRF) funded by the Ministry of Education (2017R1C1B2004616) and Cooperative Research Program for Agriculture Science and Technology Development (Supportive managing project of Center for Companion Animals Research) by Rural Development Administration (PJ0138772018).

## Supporting information

**S1 Fig. Stability assay for the survival of phage pVa-21.** Stability of phage pVa-21 was assessed at various pH (A) and temperatures (B) for 1 h.

**S2 Fig. Biofilm formation of *Vibrio sp*. used in this study.** OD_595_, optical density at 595 nm, represents biofilm forming capacity after 24 h incubation.

**S3 Fig. Plaques of pVa-21 formed in double-layer agar plates.**

**S1 Table. Host range of phage pVa-21 against all bacterial strains used in this study.**

**S2 Table. Functional categories of the predicted open reading frames (ORFs) in bacteriophage pVa-21.**

